# Ataxin-2 is essential for cytoskeletal dynamics and neurodevelopment in *Drosophila*

**DOI:** 10.1101/2021.01.07.425768

**Authors:** Urko del Castillo, Rosalind Norkett, Wen Lu, Anna Serpinskaya, Vladimir I. Gelfand

## Abstract

Ataxin-2 (Atx2) is a highly conserved RNA binding protein. Atx2 undergoes polyglutamine expansion leading to Amyotrophic Lateral Sclerosis (ALS) or Spinocerebellar Ataxia type 2 (SCA2). However, the physiological functions of Atx2 in neurons remain unknown. Here, using the powerful genetics of *Drosophila,* we show that Atx2 is essential for normal neuronal cytoskeletal dynamics and organelle trafficking. Upon neuron-specific Atx2 loss, the microtubule and actin networks were abnormally stabilized and cargo transport was drastically inhibited. Depletion of Atx2 caused multiple morphological defects in the nervous system of 3^rd^ instar larvae. These include reduced brain size, impaired optic lobe innervation and decreased dendrite outgrowth. Defects in the nervous system caused loss of the ability to crawl and lethality at the pupal stage. Taken together, these data mark Atx2 as a major regulator of cytoskeletal dynamics and denote Atx2 as an essential gene in neurodevelopment, as well as a neurodegenerative factor.

## Introduction

Ataxin-2 (Atx2) is a highly conserved RNA binding protein which is known to associate with polyribosomes, coordinate RNA granule assembly and regulate translation for synaptic plasticity (Bakthavachalu et al., 2018; McCann et al., 2011; Satterfield et al., 2002; Sudhakaran et al., 2014). Atx2 contains mRNA binding domains, intrinsically disordered domains and a polyQ domain, shown to undergo expansion, leading to neurodegenerative diseases Spinocerebellar Ataxia type 2 (SCA2) and Amyotrophic Lateral Sclerosis (ALS) (Elden et al., 2010; Riess et al., 1997). Due to its causative role in neurodegenerative diseases, there has been much interest in discerning the function of Atx2. However, whilst Atx2 is highly expressed in the developing central nervous system, we presently do not know how Atx2 functions in normal neurodevelopment. In one study, Atx2 knockout mice were generated (Kiehl et al., 2006). These mice displayed no obvious abnormalities, likely due to compensation by a related protein called Atx2-like. Therefore, the physiological roles of Atx2 remain undetermined. The gene encoding Atx2-like is absent in *Drosophila*, marking them as an ideal model organism to probe physiological Atx2 function.

Atx2 can associate with its target RNAs, and stabilizes these transcripts (Satterfield and Pallanck, 2006; Singh et al., 2020). In turn, this action contributes to long term habituation or memory formation (Bakthavachalu et al., 2018; Sudhakaran et al., 2014). Via RNA organization and control of translation, Atx2 may be a critical regulator in many cellular processes, among them cytoskeletal dynamics. For example, Atx2 has been previously indicated in actin regulation in the germline (Satterfield et al., 2002) and microtubule organization in mitosis (Gnazzo et al., 2016; Stubenvoll et al., 2016). Collectively, these observations are of note because there are strong parallels between organization of the cytoskeleton in mitosis and in neurons (Del Castillo et al., 2020; Cheerambathur et al., 2019; Hertzler et al., 2020; Lin et al., 2012; Norkett et al., 2020; Zhao et al., 2019). In mitosis, the cytoskeleton must be reorganized to facilitate chromosome separation and actin ring contraction (Basant and Glotzer, 2018; Mishima et al., 2002). In neurons, development of neurites is highly dependent upon dynamics of the actin and microtubule cytoskeleton (Lu et al., 2013; Papandréou and Leterrier, 2018; Roossien et al., 2014).

We used the powerful genetics of *Drosophila* to probe the physiological roles of Atx2. Using this system, we show that loss of Atx2 at early developmental stages leads to ‘hyperstable’ microtubule and actin networks, and severely reduced organelle transport. Subsequently we show that neurodevelopment is drastically impaired and that loss of Atx2 in the nervous system is 100% lethal at the pupae stage.

## Results

### Ataxin-2 depletion suppresses microtubule and actin dynamics

Previous studies have shown an essential role for Atx2 in cell division via regulation of centrosomes and the spindle midzone (Gnazzo et al., 2016; Stubenvoll et al., 2016). These structures are key microtubule organizing and containing structures respectively. Further, it is well documented that mitotic machinery is repurposed in neurons to facilitate cytoskeleton reorganization (Baas, 1999; Del Castillo et al., 2019). Whilst Atx2 has known roles in neurodegeneration and is highly expressed in the developing nervous system, its physiological role in neurons remains undetermined. Therefore, we hypothesized that Atx2 may be a key player in neuronal cytoskeletal dynamics at earlier developmental stages.

We chose to investigate the properties of microtubules in *Drosophila* neurons and S2 cells – an efficient system to study the cytoskeleton using dsRNA strategies (Jolly et al., 2010). We depleted Atx2 in interphase *Drosophila* S2 cells and carried out immunostaining for acetylated tubulin. Tubulin acetylation increases with microtubule age, so is a read out for microtubule lifetime (Janke and Magiera, 2020; Portran et al., 2017; Szyk et al., 2014). In S2 cells, we found a roughly 3-fold increase in acetylated tubulin signal upon Atx2 depletion (Fig. 1A, B). In order to study the physiological role of Atx2 in neurons, we depleted the protein by neuron specific expression of shRNA against Atx2 using the pan-neuronal driver elav-Gal4. Crucially, this is a post-mitotic driver, so any role for Atx2 in cell division is not affected. We prepared primary neuronal cultures of *Drosophila* 3^rd^ instar larvae and found acetylated tubulin signal increased around 30% in Atx2 RNAi neurons compared to control (Fig. 1C, D). Atx2 knockdown was confirmed by western blot of brain lysate (Fig. S1). Further, we investigated acetylated tubulin levels in brain lysate from elav>Atx2 RNAi larvae brains. Consistent with our immunostaining data, western blotting demonstrated an increase in acetylated tubulin in Atx2 RNAi brain lysate compared to control (Fig. 1E, expanded in Fig. S1B). Taken together, these results suggest an increase in microtubule stability upon loss of Atx2 and confirm this effect is conserved between dividing and post mitotic cells.

**Figure 1:**
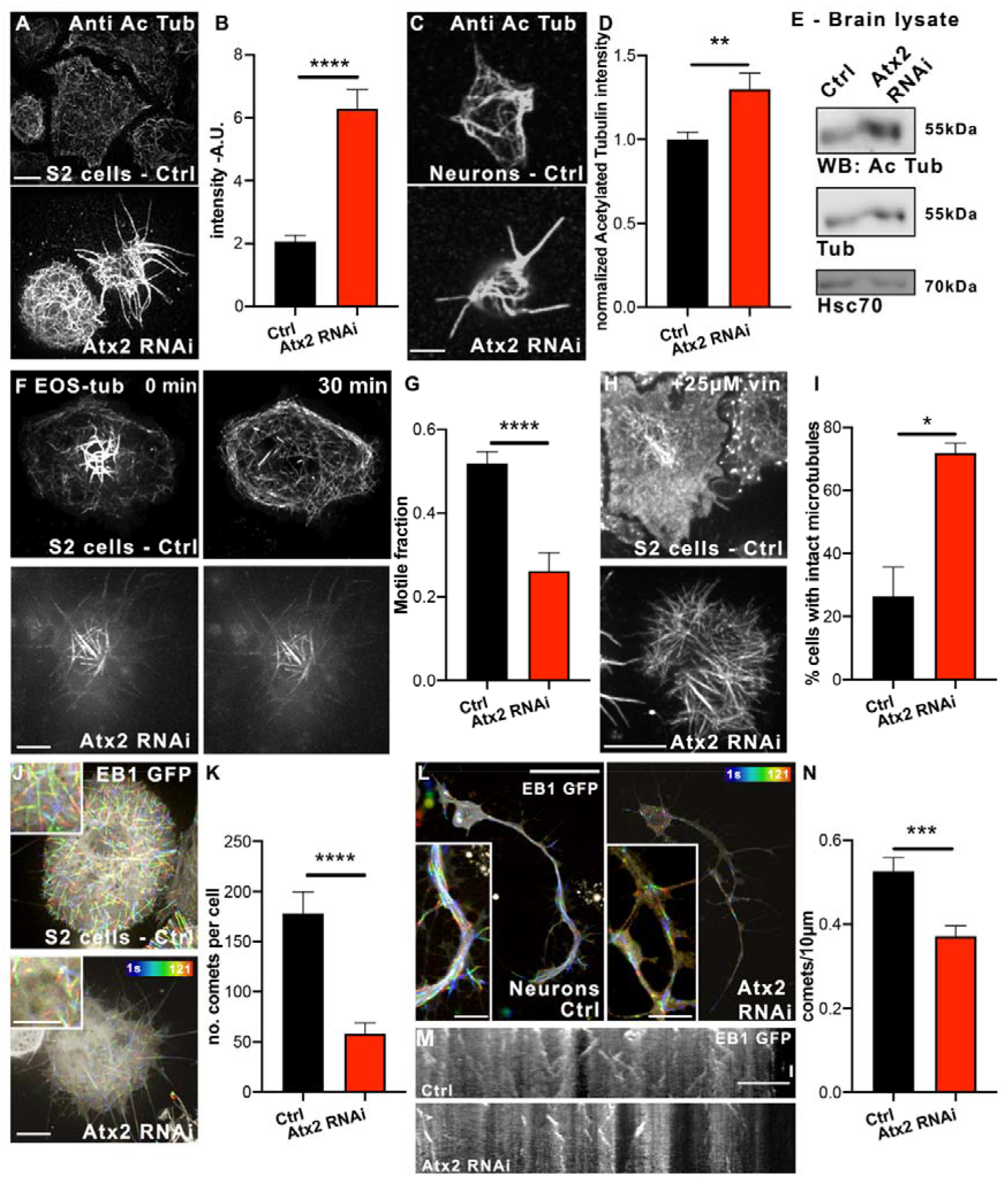
Ataxin-2 depletion suppresses microtubule dynamics. A. Example images showing Atx2 depletion in *Drosophila* S2 cells increases microtubule acetylation as quantified in B (Signal intensity Control = 2061210 ± 204596 A.U, n = 50 cells. Atx2 RNAi = 6286404 ± 608574, n = 41 cells) C. Example images showing Atx2 depletion increases microtubule acetylation in *Drosophila* neurons in culture (scale bar, 10μm), as quantified in D (Signal Intensity Control = 1.0 ± 0.04, n = 23 cells, Signal Intensity Atx2 RNAi = 1.3 ± 0.10, n = 27 cells, p = 0.01). E. Western blot of acetylated tubulin levels in Control and elav>Atx2 RNAi brain lysate from *Drosophila* 3^rd^ instar larvae. F. Example images of EOS-tubulin in S2 cells 0 and 30mins after photoconversion (scale bar, 10μm) as quantified in G (Motile fraction Control = 0.52 ± 0.03, n = 21 cells, Atx2 RNAi = 0.26 ± 0.04, n = 22 cells, p < 0.0001). H. Example images of tubulin staining in S2 cells showing microtubules after 25μM vinblastine treatment for 1hr. Scale bar, 10μm. I. Quantification of cells with an intact microtubule network after vinblastine treatment (Control = 26.4% ± 9.4, n = 38 cells, Atx2 RNAi = 71.9% ± 3.1, n = 22 cells, p = 0.04). J. Temporal color code images showing motility of EB1-GFP comets in S2 cells over 120 seconds. Scale bar, 10μm. K. Quantification of number of EB1 comets per cell (Control = 178.2 ± 21.5 comets, n = 17 cells, Atx2 RNAi = 58.0 ± 10.7, n = 15 cells, p < 0.001). L. Temporal color code images showing motility of EB1-GFP comets in cultured neurons over 120 seconds. Scale bar, 10μm, inset, 5μm. M. Kymographs showing motility of EB1 comets in neuronal processes over 120 seconds. Scale bar, 10μm. N. Quantification of number of EB1 comets per process length (Control = 0.53 ± 0.03, n=31, Atx2 RNAi = 0.37 ± 0.02, n=34, p = 0.0004).

To investigate whether the increase in acetylated tubulin was related to an alteration in microtubule dynamics, we directly measured microtubule dynamics using *Drosophila* S2 cells expressing photoconvertible EOS-tubulin. UV-light was applied to a specific region to convert EOS-tubulin fluorescence from green to red and the photoconverted red signal outside the original conversion region was quantified 30 minutes after photoconversion. Incorporation of photoconverted tubulin subunits to microtubules located outside the original photoconverted area directly demonstrates tubulin subunit exchange. Atx2 depletion led to around a 50% decrease in photoconverted signal outside the ROI (Fig. 1F, G, movie 1). This indicates that Atx2 is necessary for normal subunit exchange. Additionally, we investigated the stability of microtubules after treatment with a microtubule depolymerizing agent vinblastine by immunostaining for tubulin. Consistent with our previous data, we found that Atx2 depletion confers resistance of microtubules to vinblastine – intact microtubules could be detected in the Atx2 RNAi cells, whereas microtubules in the control cells were largely depolymerized (Fig. 1H and I).

In order to directly assess the effect of loss of Atx2 on microtubule dynamics, we carried out live imaging of EB1 (end binding protein 1) – a marker for polymerizing microtubule plus-ends (Akhmanova and Steinmetz, 2015). Initially, we observed EB1-GFP comets in S2 cells (Fig. 1J and K, movie 2). We found around a 65% decrease in the number of comets per cell, as indicated by the loss in comets shown in the temporal color code – a color coded projection of each comet over time. Furthermore, we carried out the same assay in cultured larvae neurons expressing EB1-GFP (Fig. 1L, movie 3). Comets in neuronal processes are demonstrated in kymographs in Fig. 1M where time is projected on the Y axis, showing motile objects as diagonal lines. Upon Atx2 knockdown, we found a roughly 30% decrease in the number of comets per 10μm (Fig. 1N). Notably, the total microtubule content in these cells is unchanged, as shown by tubulin staining of extracted control and Atx2 knockdown neurons (Fig. S2A, B). Therefore, Atx2 knockdown decreases only the dynamic microtubule population, not the total amount of microtubules. Taken together, these data show that loss of Atx2 confers increased microtubule stability and that Atx2 is essential for normal microtubule dynamics, both in cultured S2 cells, and crucially, in neurons.

In addition to the microtubule cytoskeleton, Atx2 has been linked to the actin filament formation in *Drosophila* embryos (Satterfield et al., 2002). Therefore, we hypothesized that Atx2 depletion may impact actin dynamics in the nervous system. Initially, we knocked down Atx2 by RNAi in *Drosophila* S2 cells and stained for actin filaments using Rhodamine-phalloidin. We treated these cells with actin depolymerizing agent Latrunculin B (LatB). We found that in control cells, the F-actin signal, detected by phalloidin staining, was decreased by around 35%, indicative of actin depolymerization. However, upon Atx2 knock down, there was no significant alteration in F-actin signal (Fig. 2A, B). The sensitivity of F-actin to depolymerizing agent LatB decreased.

**Figure 2:**
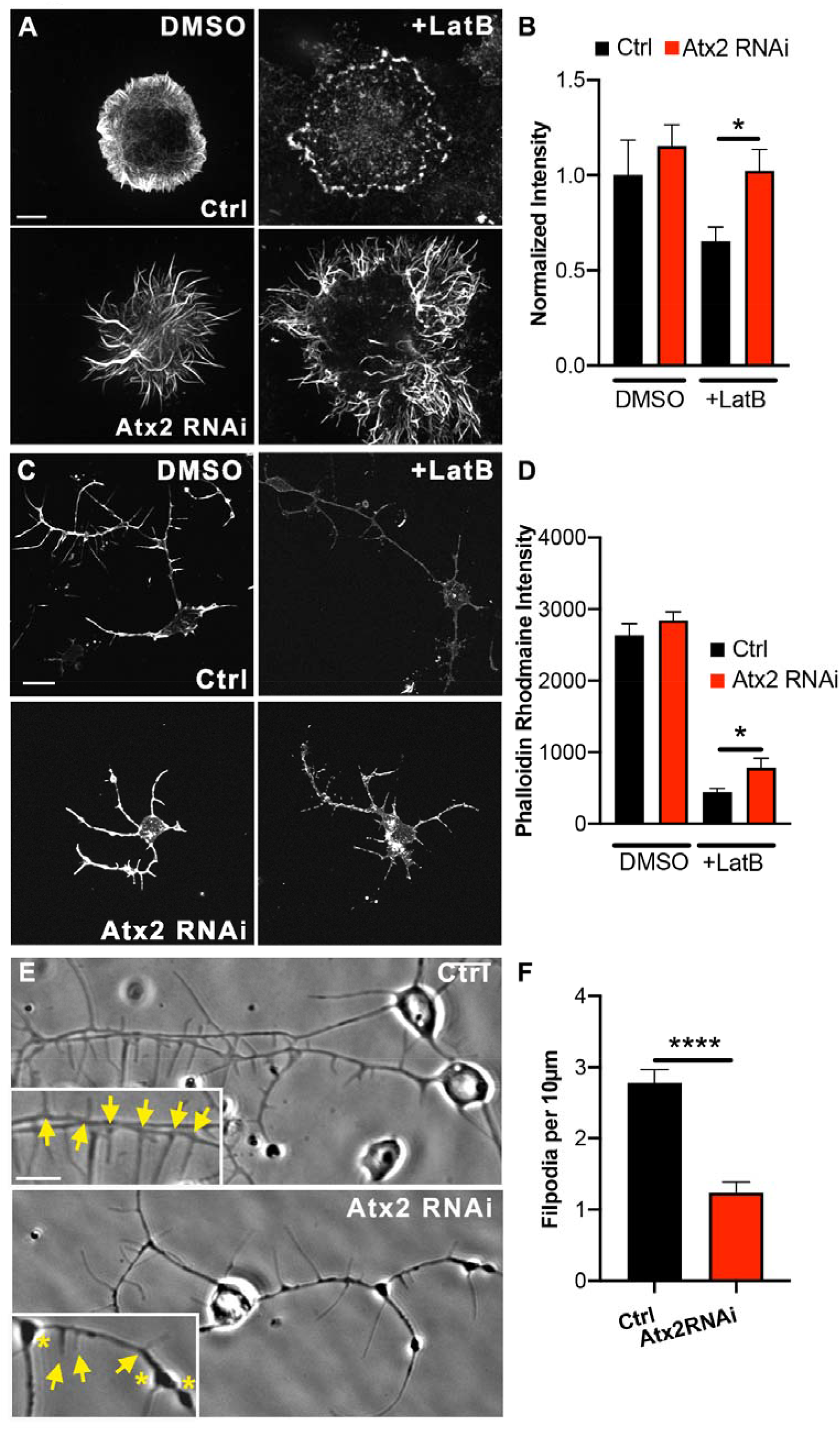
Ataxin-2 depletion stabilizes the actin cytoskeleton A. Example images showing phalloidin staining for F-actin in control and Atx2 RNAi treated *Drosophila* S2 cells treated with DMSO or 10μM LatB for 1hr. Scale bar, 10μm B. Normalized Phalloidin Rhodamine signal (Control = 1.00 ± 0.19, Atx2 RNAi = 1.15 ± 0.11, Control + LatB = 0.65 ± 0.07, Atx2 RNAi + LatB = 1.02 ± 0.11, n = 15-24 cells, p = 0.03). C. Example images showing phalloidin staining in control and elav>Atx2 RNAi neurons treated with DMSO or 10μM LatB for 1hr. Scale bar, 10μm. D. Quantification of Phalloidin Rhodamine intensity along length of neuron (Signal Intensity Control LatB = 440.4 ± 53, n = 37 cells, Signal Intensity Atx2 RNAi LatB = 786.5 ± 128, n = 27 cells, p = 0.02). E. Example phase contrast images showing filopodia from neuronal processes in control and elav>Atx2 RNAi neurons. Scale bar, 10μm. F. Average number of filopodia per 10μm process length (Control = 2.8 ± 0.19, n = 28 cells, Atx2 RNAi = 1.2 ± 0.15, n = 28 cells, p <0.0001).

To test this actin regulation in neurons, we carried out similar experiments in cultured neurons from *Drosophila* 3^rd^ instar larvae, either wild type, or after Atx2 knock-down. We demonstrated that similarly to S2 cells, actin filaments in neurons are more stable after Atx2 knockdown. As demonstrated by Rho-phalloidin staining about 40% more F-actin survived the treatment with LatB (Fig. 2C, D). This shows that actin polymers are less sensitive to depolymerizing agents upon loss of Atx2, and so, that the actin network is less dynamic. Therefore, Atx2 is an essential actin regulator in dividing and non-dividing cells.

In order to study the potential effects of increased actin stability on neuronal morphology, we chose to observe filopodia, thin rod like protrusions rich in F-actin, essential for generation of neurites and the growth cone (Lin et al., 1996). We prepared primary neuronal cultures from 3^rd^ instar larvae and carried out phase contrast imaging to examine filopodia along neuronal processes. Upon loss of Atx2, the number of filopodia projecting from neurites decreased over 50% (Fig. 2E, F, see arrows). This is consistent with a less dynamic actin network upon loss of Atx2, with filopodia being unable to form.

These data show that Atx2 is essential for normal actin dynamics in neurons. These effects are comparable to our data demonstrating the microtubule network is stabilized upon Atx2 knockdown, and highlight the role of Atx2 in regulating normal cytoskeletal dynamics in neurons during development of the nervous system.

### Ataxin-2 is essential for organelle transport and distribution in *Drosophila* neurons

Beyond neuronal morphology, both actin and microtubule networks are crucial for normal organelle transport and distribution. This is essential in neurons due to their unique architecture. Microtubules act as tracks for longer range transport of cargo by kinesin and dynein motors, whereas actin is responsible for shorter range transport and cargo anchoring by myosin motors (Kapitein et al., 2013; Lu et al., 2020; Noordstra et al., 2016; Twelvetrees, 2020; De Vos and Hafezparast, 2017). Therefore, we hypothesized that the effects of loss of Atx2 on the cytoskeleton would have a crucial impact on organelle trafficking.

To investigate the effect of loss of Atx2 on organelle trafficking, we started by carrying out live imaging of two cargoes in *Drosophila* 3^rd^ instar larvae neurons in culture – mitochondria and lysosomes (Fig. 3A). Using the neuron specific driver elav-Gal4, we labelled mitochondria using a genetically encoded mCherry fluorescent protein targeted to the outer mitochondrial membrane. Time lapse microscopy showed a 60% reduction in motile mitochondria. This is exemplified in the kymographs in Fig. 3B and in movie 4 and quantified in Fig. 3D. We also studied the motility of lysosomes labeled with lysotracker. Consistent with our data for mitochondrial transport, we found a vast decrease in lysosome motility upon loss of Atx2 (Fig. 3C, movie 5) – in control conditions around 30% of the organelles were moving, this dropped about two-fold in Atx2 RNAi neurons (Fig. 3E).

**Figure 3:**
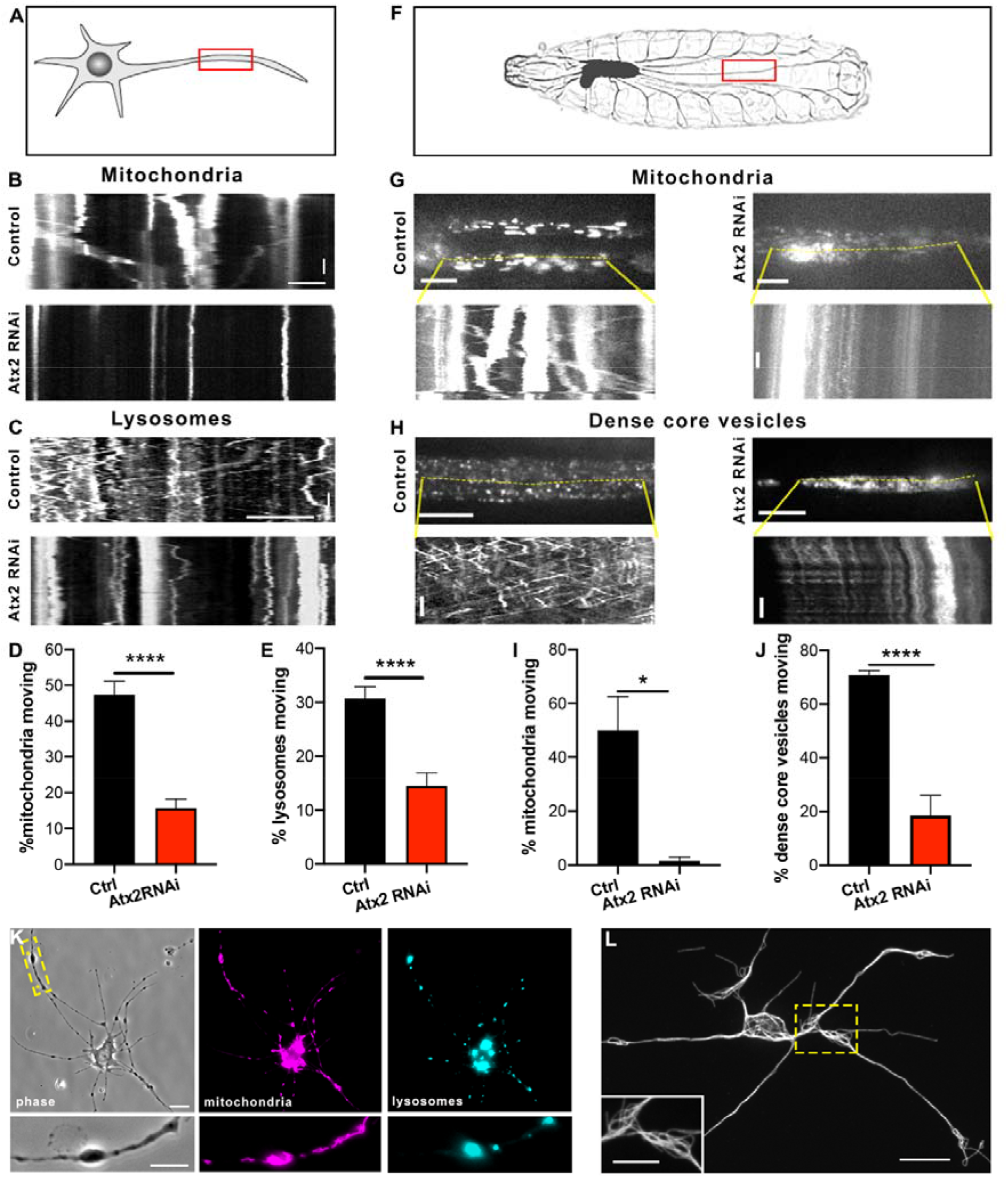
Ataxin-2 depletion inhibits organelle transport A. Schematic showing region of imaging in neurons in culture. B. Example kymographs showing mitochondria motility in processes of control and elav>Atx2 RNAi neurons. Horizontal scale bar, 10μm. Vertical scale bar, 20s. C. Example kymographs showing lysosome motility in processes of control and Atx2 RNAi neurons. Horizontal scale bar, 10μm. Vertical scale bar, 20s. D. Quantification of motile mitochondria in control and Atx2 knockdown neurons (Control = 47% ± 3.7, n = 25 cells, Atx2 RNAi = 15.7% ± 2.5, n = 28 cells, p <0.0001). E. Quantification of motile lysosomes in control and Atx2 knockdown neurons (Control = 31% ± 2.1, n = 25 cells, Atx2 RNAi = 14.5% ± 2.4, n = 29 cells, p <0.0001). F. Schematic image of a third instar larva showing segmental nerves. Example kymographs showing mitochondria motility (G) and dense core vesicle motility (H) in segmental nerves *in vivo* in control and elav>Atx2 RNAi larvae. Vertical scale bars, 10s, horizontal scale bars,10μm for dense core vesicles, 20μm for mitochondria. I. Quantification of motile mitochondria in control and Atx2 knockdown segmental nerves *in vivo* (Control = 50% ± 12.5, n = 3 larvae, Atx2 RNAi = 1.5% ± 1.5, n = 3 larvae, p = 0.015). J. Quantification of motile DCVs in control and Atx2 knockdown segmental nerves *in vivo* (Control = 66% ± 1.7, n = 9 larvae, Atx2 RNAi = 18.6% ± 7.4, n = 8 larvae, p < 0.0001). K. Example image showing mitochondria and lysosome distribution in an elav>Atx2 RNAi cultured neuron. Scale bar, 10μm, inset 5μm. L. Example super resolution images of microtubules in varicosities in Atx2 RNAi neurons. Scale bar 10μm, inset 5μm.

Next, we extended these findings to investigate organelle transport *in vivo* in the segmental nerves of *Drosophila* 3^rd^ instar larvae. These nerves contain the axons of motor and sensory neurons which extend from the CNS into segments of the larvae body wall (Fig. 3F). We studied motility of two cargoes; in this case mitochondria and dense core vesicles (presynaptic carriers). As in culture, we found that *in vivo* mitochondrial transport was drastically decreased by around 90% in Atx2 RNAi animals compared to control (Fig. 3G, I, movie 6). Trafficking of dense core vesicles was similarly impaired in Atx2 RNAi larvae – motile organelles decreased by around 70% (Fig. 3H, J, movie 7). Taken together these data show Atx2 has an essential role in normal transport of multiple organelles transport in neurons *in vitro* and *in vivo*.

In our live imaging experiments, we noted accumulations of stationary organelles (e.g. movie 6) and we have previously shown that depleting Atx2 leads to varicosities in neuronal processes containing ER accumulations (Fig. 2E asterisks) (del Castillo et al., 2019). Downstream of the Atx2 dependent loss of cargo transport, we investigated the distribution of organelles throughout neurons in culture. Phase contrast imaging coupled with labelling of mitochondria and lysosomes demonstrated that these organelles could be found colocalized in the axonal varicosities (Fig. 3K). Indeed, the majority of the varicosities held both of these organelles (84%) while only around 5% each of varicosities contained only one or neither of these organelles. This is consistent with a ‘traffic jam’ leading to accumulation of organelles at these sites due to severely impaired transport.

We were interested in the microtubule structure at these ‘traffic jam’ sites. In order to visualize microtubules at the sites, we carried out super resolution imaging on extracted Atx2 RNAi neurons with stained for tubulin. We found that microtubules in these regions are not organized in parallel arrays, but instead were disordered and looped (Fig. 3L). This disorganization of the microtubule network may prevent normal organelle transport through these sites, causing their accumulation.

The cytoskeletal and trafficking phenotypes we observe are likely dependent on the RNA transcripts regulated by Atx2. We carried out RNAseq analyses of dissected control vs Atx2 RNAi 3^rd^ instar larvae brains to identify differentially expressed genes that may regulate the cytoskeleton. Depletion of Atx2 induced clear alterations in the neuronal transcriptome. We identified 1636 significantly upregulated and 1684 significantly downregulated transcripts out of 16729 tested (adjusted p value < 0.05) (Fig. S3A). This data set confirms a decrease in the level of Atx2 mRNA in the Atx2 knockdown condition compared to control (Log_2_FC −0.97, equivalent to around a 50% decrease). Next we grouped the differentially expressed genes according to their indexed cellular component using PANTHER gene ontology analysis (Thomas et al., 2003). The number of transcripts per group in the data set is compared to the expected number, based on the total number of transcripts in this group in the *Drosophila* reference transcriptome. Therefore, any ‘overrepresented’ cellular components can be identified according to the list of differentially expressed genes. Amongst enriched cellular components, cytoskeletal proteins were around 1.5 fold overrepresented (GO:0005856, p = 4.73E-03), and components of the actin cytoskeleton are around 2 fold overrepresented (GO:0015629, p = 5.19E-03) (Fig. S3B). We further probed these two groups to identify the protein class or molecular function assigned to these transcripts and found 50% of these were ‘classical’ cytoskeletal proteins (rather than protein modifiers), which we define as proteins annotated ‘PC00085’ in the database (Table S1). Therefore, Atx2 has the ability to regulate mRNA levels of many cytoskeleton components.

### Ataxin-2 is essential for neurodevelopment, locomotion and viability in *Drosophila*

We and others have demonstrated that the cytoskeleton is one of the major drivers of neurite initiation and neuronal development (Del Castillo et al., 2015; Korobova and Svitkina, 2008; Lu et al., 2013; Norkett et al., 2020; Roossien et al., 2014; Winding et al., 2016; Yu et al., 2000). Therefore, we studied how the nervous system develops in the absence of Atx2 as a result of severely impaired cytoskeletal dynamics and trafficking. Firstly, we studied the size of the brain in intact Drosophila 3^rd^ instar larvae. We used the gene trap line Nrv2 GFP to endogenously label the central nervous system and segmental nerves (Fig. 4A). We found the length of the ventral nerve cord (VNC), normalized to body length, was decreased by around a third (Fig. 4B, C). The length of the VNC was decreased whilst the length of the larva body was unchanged between control and Atx2 RNAi animals (Fig. S4A, B). Therefore, Atx2 is essential for correct brain development.

**Figure 4:**
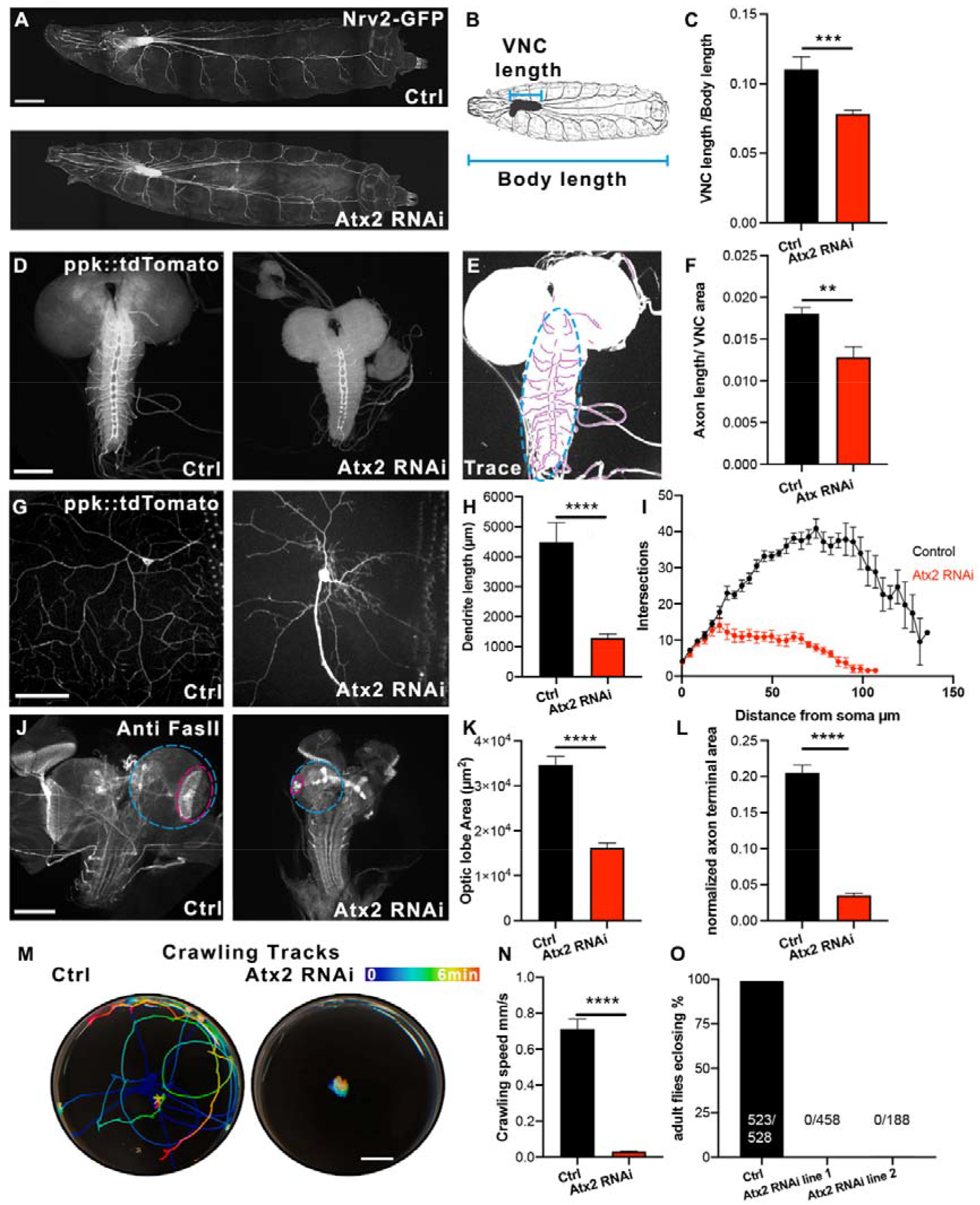
Ataxin-2 is essential for neurodevelopment, locomotion and viability in *Drosophila.* A. Atx2 depletion decreases brain size. Example images of control and elav>Atx2 RNAi brains from *Drosophila* 3^rd^ instar larvae. Scale bar 100μm B. Illustration showing brain length and body length measurements C. Length of VNC normalized to body length. (Control ratio = 0.11 ± 0.009, Atx2 RNAi ratio = 0.08 ± 0.003, n = 8 control and 10 Atx2 RNAi animals, p = 0.0006). D. Atx2 depletion decreases axon development. Example images of control and elav>Atx2 RNAi brains from *Drosophila* 3^rd^ instar larvae. E. Illustration showing example of axon tracing (magenta) in the VNC (cyan). F. Axon length in VNC normalized to VNC area (Control = 0.018 ± 7×10^−4^, Atx2 RNAi ratio = 0.013 ± 1.3×10^−3^, n = 8 control and 10 Atx2 RNAi animals, p = 0.006). G. Atx2 depletion decreases dendrite development. Example images of DA neurons in control and elav>Atx2 RNAi neurons. Scale bar, 50μm. H. Dendrite length per cell in control and elav>Atx2 RNAi neurons (Control = 4500 μm ± 644, Atx2 RNAi = 1290 μm ± 134, n=11 control and 11 Atx2 RNAi animals, p < 0.0001). I. Sholl analysis of control and elav>Atx2 RNAi neurons DA neurons showing number of intersections with distance from soma (n=6 control and 4 Atx2 RNAi animals). J. Atx2 depletion decreases optic lobe innervation. Example images of control and elav>Atx2 RNAi brains from *Drosophila* 3^rd^ instar larvae. Scale bar, 100μm. K. Size of optic lobes in control and elav>Atx2 RNAi brains (Control = 34662 μm^2^ ± 1897, Atx2 RNAi = 16198 μm^2^ ± 1128, n=23 control and 20 Atx2 RNAi brains, p<0.0001). L. Area of optic stalks normalized to optic lobes in control and Atx2 RNAi brains (Control = 0.205 ± 0.01, Atx2 RNAi = 0.035 ± 0.003, n=23 control and 20 Atx2 RNAi brains, p<0.0001). M. Atx2 depletion impairs locomotion. Example crawling tracks of control and elav>Atx2 RNAi 3^rd^ instar larvae. Scale bar = 10mm. N. Crawling velocity of control and elav>Atx2 RNAi 3^rd^ instar larvae (Control = 0.71 mm/s ± 0.05, Atx2 RNAi = 0.03 mm/s ± 0.003, n = 12 control and 12 Atx2 RNAi animals). O. Atx2 depletion is lethal at the pupae stage. Percentage of adults eclosing from pupae cases (Control = 99%, Atx2 RNAi line 1 = 0.0%, Atx2 RNAi line 2 = 0.0%).

We investigated the effect of the loss of Atx2 on axonal and dendritic development. To do this, we labelled Class IV DA (Dendritic arborization) neurons (sensory neurons that line the *Drosophila* body wall) with a fluorescent reporter expressed specifically in these cells (ppk::tdTomato). To assess axon development, we dissected brains of these larvae and imaged the axonal projections of these neurons extending into the VNC (Fig. 4D) We traced the axons in this region and found the axon length, normalized to the area of the VNC, was significantly decreased upon Atx2 depletion (Illustrated in Fig. 4E, quantified in Fig. 4F). Regarding dendritic development; under control conditions, these neurons extended long, branched dendritic arbors which covered the whole body segment. However, upon loss of Atx2, we found that these dendritic arbors were drastically shrunken (Fig. 4G). This effect was quantified in two ways, firstly the total dendrite length per cell was decreased by around 65% (Fig. 4H), consistent with our previously published *in vitro* data (del Castillo et al., 2019). Secondly, to further describe the effect of Atx2 depletion on the shape of the dendritic arbor, we carried out Sholl analysis (Sholl, 1953). Concentric circles are drawn outwards from the soma, and number of intersections that the dendritic arbor makes with each circle is counted as a representation of dendrite morphology. From 20μm from the soma, the Atx2 RNAi neurons displayed many fewer intersections than control neurons and the dendritic arbors only extended to around 100μm from the soma, whereas the dendrites of the control neurons extended around 30% more than this (Fig. 4I). Therefore, Atx2 knock down severely impairs dendrite development as well as axon development. We also studied axon development in the central nervous system by dissecting 3^rd^ instar larvae brains and immunostaining for Fasciclin II (FasII) – an axon specific marker. We once more found that Atx2 RNAi brains were drastically smaller than controls. Quantification of the optic lobes (Fig. 4J, cyan) showed a roughly 2-fold decrease in area (Fig. 4K). Further we quantified area of the retinotopic pattern – the structure representing the photoreceptor axons (Fig. 4J, magenta). When normalized to the area of the optic lobe, we found this structure was decreased to around 10% of control area upon Atx2 depletion (Fig. 4L). These phenotypes demonstrate that Atx2 depletion significantly impairs axon targeting and innervation. These data are consistent with our observations that depleting Atx2 has severe consequences on cytoskeletal dynamics and organelle transport.

We studied the effect of this underdeveloped nervous system on the whole organism, initially by studying locomotion. We imaged 3^rd^ instar larvae crawling on agar plates and calculated the velocity. Control larvae crawled over the surface of the plate at about 0.7mm/s. However, Atx2 knockdown larvae were largely stationary, crawling with a velocity of less than 0.1mm/s (Movie 8). This is further shown by the color-coded tracks of each larva over time (Fig. 4M quantified in Fig. 4N).

Crucially, we found that depleting Atx2 in neurons was lethal at the pupae stage. Animals expressing the Atx2 RNAi were not viable to adult stages (Fig. 4O, control; 523/528, RNAi line 1; 0/458, RNAi line 2; 0/188 flies eclosed). There was no difference in survival between RNAi stock lines (96% viable) and controls (98% viable) when the Atx2 RNAi expression was not driven by a Gal4. We found consistent results with two different Atx2 RNAi lines; therefore, this effect is specific to Atx2 knockdown rather than an off-target effect. Taken together, these data show that Atx2 is necessary for normal development of both axons and dendrites. As a result of the loss of Atx2, development of the nervous system is severely impaired. Subsequently, the larvae are unable to move and die during development. These data highlight the role for Atx2 as an essential gene in nervous system development and that it is absolutely required for normal cytoskeletal dynamics and cargo transport.

## Discussion

Multiple works have demonstrated the involvement of mutant Atx2 in neurodegenerative conditions ALS and SCA2 (Elden et al., 2010; Riess et al., 1997). However, an understanding of the physiological roles of the RNA-binding protein in neurons is still lacking, potentially due to redundancy between Atx2 and Atx2-like in mammals. Here we use *Drosophila* as a model system to uncover the necessity of wild type Atx2 for the neuronal cytoskeleton and organelle trafficking. Further to this, we show gross morphological defects in the nervous system. These neurodevelopmental defects lead to decreased locomotion of *Drosophila* larvae and are lethal during development, highlighting the necessity of Atx2 during development.

We show that Atx2 is essential for normal cytoskeletal dynamics, in both microtubule and actin networks. Firstly, our data show that targeted depletion of Atx2 in neurons drives an increase in microtubule acetylation, a decrease in tubulin subunit exchange and fewer EB1 comets. Thus, we conclude microtubules shift to a more stable, less dynamic state. A role for Atx2 in regulating the microtubule network is consistent with previous reports of Atx2 having roles in centrosome organization and mitotic spindle regulation in *C. elegans* (Gnazzo et al., 2016; Stubenvoll et al., 2016). Secondly, we report an increase in the stability of the actin network upon Atx2 knockdown. These build upon early descriptions of Atx2 function, which showed evidence of disorganized actin bundles in retinal precursor cells, the female germline and bristles of Atx2 null *Drosophila* (Satterfield et al., 2002). Our data extend previous findings by highlighting specific neuronal effects of Atx2.

Beyond the altered cytoskeletal network, we show that decreasing levels of Atx2 decreases organelle transport in culture and *in vivo*. Impaired organelle transport is likely a direct consequence of the cytoskeletal defects caused by Atx2. Microtubule stability and longevity would be thought to facilitate microtubule motor-based organelle transport (Godena et al., 2014; Kaul et al., 2014). However, we see a decrease in motility of multiple cargoes. Instead, an increase in the stability of the actin network may lead to organelles being anchored or trapped by myosin motors (Kapitein et al., 2013; Lu et al., 2020). Alternatively, changes in mRNA levels of motor proteins may contribute to decreased trafficking (table S1), and would explain long range transport defects rather than organelles only stationary in varicosities. However, the changes in transcript levels for these proteins are modest.

This loss of organelle transport leads to ‘traffic jams’, accumulations of organelles in axonal varicosities. These axonal varicosities are also known to contain ER aggregates (del Castillo et al., 2019). As well as essential for development, improper organelle transport can be seen in neurodegenerative diseases, including mitochondria trapped in dystrophic neurites in an Alzheimer’s disease model (Correia et al., 2016; Fiala et al., 2007), and impairments in endosome retrograde transport in ALS (Xie et al., 2015).

Finally, we demonstrate the necessity of wild-type Atx2 for normal neurodevelopment. This advances our understanding of Atx2 function beyond the information provided from the knock out mouse model by overcoming redundancy between mammalian Atx2 and Atx2-like, and by highlighting nervous system specific effects of Atx2 loss. Neuron specific depletion of Atx2 led to loss of normal axon and dendrite formation in the central and peripheral nervous systems, smaller brains normalized to body size, and, ultimately locomotion deficits and death at the pupae stage. This is significant as Atx2 is typically studied as a neurodegenerative risk factor. However, our data highlight its physiological role at earlier developmental stages, yet independently of any roles in mitosis. Microtubule and actin dynamics are intricately linked with neurite outgrowth. Microtubules are crucial to transduce forces that are generated by molecular motors, which control neuronal polarization and morphology (Del Castillo et al., 2019; Lin et al., 2012; Lu et al., 2013; Norkett et al., 2020; Roossien et al., 2014; Winding et al., 2016; Zheng et al., 2008). In neurons in particular, actin dynamics have important consequences on development via growth cone formation and normal axon outgrowth (Lin et al., 1996; Papandréou and Leterrier, 2018). Incorrectly regulated actin dynamics may also impair the organization of the microtubule cytoskeleton. Actin is necessary to guide microtubules in the axon, at filipodia and at the growth cone (Bradke and Dotti, 1999; Hahn et al., 2020; Korobova and Svitkina, 2008; Schaefer et al., 2008). Therefore, impaired microtubule and actin dynamics may contribute to the gross neurite extension defects we describe. Furthermore, correct cargo trafficking and delivery, mediated by microtubule tracks and the actin network, is essential for axon development and dendritic arborization (Guedes-Dias and Holzbaur, 2019; Norkett et al., 2016; Zheng et al., 2008). In turn, this impairment in organelle distribution could lead to the neuronal defects we observed with loss of Atx2 at early developmental stages.

It will be crucial to specify the mechanisms by which Atx2 can act as a major regulator of the neuronal cytoskeleton. Whilst the mechanism is likely via translational control, protein-protein interaction remains a possibility. It is known that Atx2 does not associate directly with actin (Satterfield et al., 2002), yet Atx2 does associate with actin regulator profilin, another ALS risk factor which locates to stress granules (Figley et al., 2014; Ralser et al., 2005). However, as we show here, among the target mRNAs of Atx2 are actin and microtubule cytoskeleton components and molecular motors. These data demonstrate the ability of Atx2 to regulate multiple cytoskeletal networks. Atx2 tightly regulates RNP granules and is necessary for axon path finding (Bakthavachalu et al., 2018; Singh et al., 2020). It is essential for local translation for synaptic plasticity (McCann et al., 2011). Aberrant localization and translation of these transcripts could underlie the phenotypes herein reported. Further exploration of these, and our data will be invaluable in fully understanding the physiological roles of Atx2 in the neuron.

The data we present here address the physiological roles of Atx2 via a neuron specific knockdown approach. Specifically, we study the effect of Atx2 depletion on the developing rather than mature CNS. These data complement studies of mutant, polyQ expanded Atx2 in neurodegeneration – widely believed to cause a toxic gain of function (Huynh et al., 2000). Our data are also essential to thoroughly understand the mechanism by which Atx2 knockdown ameliorates TDP-43 pathology in ALS (Becker et al., 2017). Therefore, our data are relevant to ALS disease mechanisms and therapies. Further, our use of *Drosophila* as a model organism circumvents the presence of Atx2-like/Ataxin2 related proteins in mammals. Curiously, recent reports of an Atx2-like knockout mouse demonstrate lethality at early developmental stages and impaired neural development as shown by thinner cortices (Key et al., 2020). These observations are consistent with our data, highlighting conservation of roles of Atx2 and its related proteins across taxa.

## Supporting information

resources table

## Author Contributions

UdC, RN and VG designed research, UdC, RN, WL and AS performed research, UdC and RN analyzed data, RN, UdC and VG wrote the manuscript.

## Funding sources and Acknowledgments

We thank members of the Gelfand lab and Dr. M. Gelfand for helpful discussion. Stocks obtained from the Bloomington Drosophila Stock Center (NIH P40OD018537) were used in this study. We thank Drs. A. Lim, W. Saxton and S. Rogers for *Drosophila* lines. This work was supported by the Northwestern University NUSeq Core Facility. Research reported in this study was supported by National Institute of General Medical Sciences Grants R01GM052111 and R35GM131752 (to V.I.G.).

## Star Methods

### Resource availability

#### Lead contact

Further information and requests for resources and reagents should be directed to and will be fulfilled by the lead contact, Vladimir Gelfand.

#### Materials availability

This study did not generate new unique reagents

#### Data and Code availability

RNA-seq data will be deposited at GEO to be publicly available as upon acceptance of manuscript. Microscopy data reported in this paper will be shared by the lead contact upon request.

This study did not generate any original code.

Any additional information required to reanalyze the data reported in this paper is available from the lead contact upon request.

## Materials and Methods

### Fly stocks and plasmids

Fly stocks and crosses were cultured on standard cornmeal food at room temperature based on the Bloomington Stock Center recipe. Fly stocks used in this study are listed in the key resources table. All experiments used Atx2 shRNA line BDSC #44012 unless otherwise stated. For tubulin photoconversion experiments, S2 cells were transfected with pMT-EOS-tubulin as described in Barlan et al (Barlan et al., 2013).

### *Drosophila* Cell Culture: Primary Neurons and S2 Cells

*Drosophila* S2 cells were maintained in Insect-Xpress medium (Lonza) at 25°C. These cells came directly from DGRC and their identity was not subsequently confirmed. Transfections were carried out with Effectene (Qiagen) according to the manufacturer’s instructions. dsRNA was added to cells on days 1 and 3 and imaging was carried out on day 5. For EB1 experiments a stable S2 cell line was created expressing EB1-GFP under control of the EB1 promoter in the pMT vector. Primary neuronal cultures were prepared as previously described (Norkett et al., 2020). Briefly, brains from 3rd instar larvae were dissected and tissue was dissociated using liberase (Roche). Cells were plated on ConA-coated glass coverslips and maintained in Schneiders medium supplemented with 20% FBS, 5 μg/ml insulin, 100 μg/ml Pen-Strep, 50 μg/ml Gentamycin and 10 μg/ml Tetracycline.

### Development, Locomotion and Survival analysis

To assess development of the nervous system, Atx2 RNAi was driven by elav gal4 in flies expressing endogenously GFP tagged Nrv2, labelling the CNS. Larvae for analysis were staged based on date of egg hatching and body size to ensure the same age animals were used in control and Atx2 RNAi conditions. No differences were noted between the groups for time taken for egg hatching and body size. For locomotion assays, 3^rd^ Instar larvae were allowed to crawl on the surface of a petri dish containing 2% agarose supplemented with apple juice and charcoal powder. Trajectories are presented using the Temporal-Color Code plugin (Physics LUT) in Fiji. For line 44012 (which we designated Atx2 shRNA line 1), homozygous adults were crossed with elav-gal4 flies and number of adults eclosing from pupae cases was counted. For line 36114 (designated Atx2 shRNA line 2), heterozygous adults were crossed with elav-gal4 flies and number of ‘non tubby’ adults eclosing from pupae cases. Elav-gal4 flies were used as control.

### Immunostaining, Western Blotting and Antibodies

Antibodies used in this study are described in the key resources table. Optic lobes, dissected from third-instar larva in 1 × PBS, were fixed in 4% (wt/wt) formaldehyde (methanol-free) in PBT (1 × PBS, 0.1% Triton X-100) for 20 min; brains were washed five times with PBTB (PBT + 0.2% bovine serum albumin) for 10 min and blocked in 5% (vol/vol) normal goat serum for 1 h. Samples were incubated with primary anti-Fasciclin II antibody (1:40) at 4° C overnight, washed and incubated with secondary anti-mouse for 2h at room temperature (RT). Finally, samples were washed with PBT for 10min five times before mounting.

For immunostaining of S2 cells and cultured neurons, cells were fixed 5 mins in methanol (S2 cells) or 1% glutaraldehyde and quenched in 2 mg/ml sodium borohydride (neurons). Cells were permeabilised and blocked in wash buffer (1% BSA, 0.1% Triton X-100 in TBS) for 30mins and incubated with primary antibody in wash buffer (Anti-Acetylated tubulin 1:100-1:250, DM1a 1:1000). Coverslips were washed 3x in wash buffer and incubated with secondary antibodies (1:200). For extraction, cells were incubated in extraction buffer (1% Triton X-100, 1uM taxol, 30% glycerol in BRB-80) for 2 minutes, then in extraction buffer with 1% glutaraldehyde for 2 minutes, then 1% glutaraldehyde in PBS for 3 minutes followed by reduction as above. For western blotting, dissected third instar larvae brains or S2 cell lysate were prepared in 1% SDS, boiled in laemmli buffer and analyzed by SDS-PAGE on 8% acrylamide gels. After electrophoresis, transfer onto nitrocellulose membrane was carried out and blocking was performed in 4% milk in PBS-T. Western blotting was performed using advansta western bright quantum substrate and Licor Imagequant system. To analyze acetylated tubulin levels, membranes were first incubated with anti-acetylated tubulin antibody and developed with HRP- conjugated mouse secondary. The membrane was stripped using stripping buffer (2% SDS, 0.625M Tris pH6.8, 0.8% beta-mercaptoethanol) for 30 minutes at 37°C, washed 2x 5 minutes in PBS-T and reblocked. Then membranes were incubated with anti-tubulin antibody and developed with HRP- conjugated rabbit secondary.

### Microscopy and Image Analysis

To image DA neurons or S2 cells we used a Nikon Eclipse U2000 inverted microscope equipped with a Yokogawa CSU10 spinning disk head with a perfect focus system and a 40x 1.30 or 100x 1.45 oil immersion lens. Images were acquired using Evolve EMCCD (Photometrics) controlled by Nikon Elements 4.00.07 software. For EB1 imaging in S2 cells images were acquired at 1 fps. For organelle imaging in larvae, larvae were immobilised and frame acquisition was at 3 fps (dense core vesicles) or 0.5 fps (mitochondria). Fixed and immunostained brains and neurons were imaged on a Nikon Ti2 inverted microscope equipped with Yokogawa CSU-W1 spinning disk driven by Nikon Elements 5.20 using a 20X or 100x 1.45 oil immersion lens and CMOS prime 95B sensor (Photometrics). Whole larvae were immobilized by heat treatment at 70°C for 1 minute and mounted in 100% glycerol. Samples were imaged with a 10x 0.30 dry lens and six fields of view were stitched together in Nikon Elements. For microtubule organization images, this set-up was used with a Live-SR (Gataca systems). Live imaging of lysosomes and EB1 comets in neurons was carried out at 1fps. For photoconversion experiments we applied 405 nm light from a light-emitting diode light source (89 North Heliophor) for 5 s, using an adjustable diaphragm to constrain light and illuminate only a specific region of interest. After photoconversion, images were collected every ten minutes for 30 minutes. Imaging of filopodia, organelle distribution in culture and mitochondrial trafficking was carried out on a Nikon Ti inverted microscope with a Hamamatsu Orca Flash sensor.

Image analysis was carried out in Fiji unless otherwise stated. Total neurite length in DA neurons was measured using the ‘curve tracing’ v.0.3.5 plugin for ImageJ (https://github.com/ekatrukha/CurveTrace) and Sholl analysis plugins (https://github.com/tferr/ASA/blob/Sholl_Analysis-4.0.1). Total axon length of DA neurons in the VNC was measured using the ‘curve tracing’. VNC area, length and body length were measured manually. Tubulin motile fraction was calculated as ‘photoconverted signal outside original photoconversion zone / original photoconverted signal’ at 30 mins after photoconversion. Numbers and motility of EB1 comets, mitochondria and lysosomes in culture or *in vivo* was carried out by generating kymographs and analyzed using the ‘kymobutler’ software (Jakobs et al., 2019). Motile organelles were defined as those that moved more than 2μm over the course of the movie. Comets that moved more than 1μm over the course of the movie were counted. EB1 comets in S2 cells were counted using Nikon elements analysis software (v5.20). Numbers of filopodia and organelles in varicosities were scored manually.

### RNAseq

For RNAseq analysis, libraries from control or Atx2 RNAi 3^rd^ instar larvae brains were prepared using TruSeq stranded mRNA kit (Illumina). 3 control and 3 Atx2 RNAi libraries were prepared, generating 6 samples. The stranded mRNA-seq was conducted in the Northwestern University NUSeq Core Facility. Briefly, total RNA samples were checked for quality using RINs generated from Agilent Bioanalyzer 2100. RNA quantity was determined with Qubit fluorometer. The Illumina TruSeq Stranded mRNA Library Preparation Kit was used to prepare sequencing libraries of high-quality RNA samples (RIN>7). The Kit procedure was performed without modifications. This procedure includes mRNA enrichment and fragmentation, cDNA synthesis, 3’ end adenylation, Illumina adapter ligation, library PCR amplification and validation. Illumina HiSeq 4000 NGS Sequencer was used to sequence the libraries with the production of single-end 50 bp reads. The quality of reads, in FASTQ format, was evaluated using FastQC. Reads were trimmed to remove Illumina adapters from the 3’ ends using cutadapt (Martin, 2011). Trimmed reads were aligned to the *Drosophila melanogaster* genome (BDGP6.93) using STAR (Dobin et al., 2013). Read counts for each gene were calculated using htseq-count (Anders et al., 2015) in conjunction with a gene annotation file for BDGP6.93 obtained from Ensembl (http://useast.ensembl.org/index.html). Normalization and differential expression were calculated using DESeq2 that employs the Wald test (Love et al., 2014). The cutoff for determining significantly differentially expressed genes was an FDR-adjusted p-value less than 0.05 using the Benjamini-Hochberg method.

Following identification of significantly differentially expressed genes, the PANTHER Gene ontology database was used to identify cellular components over- and under-represented in our data set based on the Drosophila transcriptome. From the cytoskeleton and actin cytoskeleton nodes, genes were further sorted according to their protein class and those ascribed ‘cytoskeleton’ (PC00085) were identified for representation of differential expression.

### Statistical Analysis

Statistical significance between two groups was determined using the unpaired, nonparametric Mann–Whitney *U* test. Data analyses were performed with Prism v6 (GraphPad Software). Statistical significance is presented as *p<0.05, **p<0.01, ***p<0.001, ****p<0.0001. Data in Figure legends are presented as mean ± standard error.

### Data availability

All data associated with this work are included in this article and the appendix.

## Movie Legends

Movie 1: Time-lapse of control (untreated cell) or Atx2 RNAi S2 cells expressing EOS-tub. Scale bar, 10μm. Related to Fig. 1F.

Movie 2: Time-lapse of control (untreated cell) or Atx2 RNAi S2 cells expressing EB1-GFP. Scale bar, 10μm. Related to Fig. 1J.

Movie 3: Time-lapse of primary neurons expressing EB1-GFP of two different genotypes. Control (elav-gal4) and Atx2 RNAi (elav>Atx2 RNAi). Scale bar, 10μm. Related to Fig. 1L.

Movie 4: Transport of mitochondria in primary cultures of control and Atx2 RNAi expressing Mito mCherry. Scale bar, 10μm. Related to Fig. 3B.

Movie 5: Transport of lysosomes in primary cultures of control and Atx2 RNAi labelled with lysotracker. Scale bar, 10μm. Related to Fig. 3C.

Movie 6: Transport of mitochondria in motor segmental nerves of control and Atx2 RNAi expressing mitoGFP. Scale bar, 10μm. Related to Fig. 3G.

Movie 7: Transport of Dense core vesicles in motor segmental nerves of control and Atx2 RNAi expressing ANF-GFP. Scale bar, 10μm. Related to Fig. 3H.

Movie 8: Crawling behavior of control (elav-gal4) and Atx2 RNAi (elav>Atx2 RNAi) *Drosophila* 3^rd^ Instar Larvae, Related to Figure 4G.

### Tables

Differentially expressed genes in Ctrl vs Atx2 RNAi S2 cells

Differentially expressed genes in Ctrl vs Atx2 RNAi 3^rd^ instar larvae brains

