## Supplementary material for "Ataxin-2 is essential for cytoskeletal dynamics and neurodevelopment in *Drosophila*": resources table

**Key Resources Table**

| REAGENT or RESOURCE | SOURCE | IDENTIFIER |
| --- | --- | --- |
| Antibodies | | |
| Anti FasII mouse monoclonal | DSHB | Monoclonal 1D4 |
| Anti Acetylated alpha tubulin (K40) mouse monoclonal | protein tech | Cat# 66200 |
| Anti Hsc 70 K-19 goat polyclonal | Santa Cruz | Cat# sc-1059 |
| Anti Dmel Atx2 rabbit polyclonal | Li Antibodies custom preparation | N/A |
| Anti alpha tubulin DM1a mouse monoclonal | Prepared in house | N/A |
| Anti tubulin rb polyclonal | Prepared in house | N/A |
| HRP-conjugated anti mouse | Jackson | RRID: AB_2340770 |
| HRP-conjugated anti rabbit | Jackson | RRID: AB_10015282 |
| HRP-conjugated anti goat | Jackson | RRID: AB_2340390 |
| Alexafluor 647 conjugated anti mouse IgG | Jackson | RRID: AB_2340862 |
| Chemicals, peptides, and recombinant proteins | | |
| Latrunculin B | Sigma Aldrich | Cat#L5288  CAS number [76343-94-7](https://www.sigmaaldrich.com/US/en/search/76343-94-7?focus=products&page=1&perPage=30&sort=relevance&term=76343-94-7&type=cas_number) |
| Vinblastine | Sigma Aldrich | Cat#V1377  CAS number 143-67-9 |
| Rhodamine conjugated Phalloidin | Thermo Fisher Scientific | Cat#R415 |
| Effectene | Qiagen | Cat#301425 |
| Critical commercial assays | | |
| TruSeq stranded mRNA kit | Illumina | Cat#20020596 |
| Experimental models: Cell lines | | |
| S2R+ | DGRC | RRID:CVCL_A5UM |
| Experimental models: Organisms/strains | | |
| *Drosophila melanogaster* Ataxin-2 RNAi TRiP line #1  (Val20, second chromosome attP40 insertion targeting CDS 561-581) TRiP.HMS02726 | Bloomington Stock Centre (BDSC) | RRID:BDSC_44102 |
| *Drosophila melanogaster* Ataxin-2 RNAi TRiP line #2 (Val20, third chromosome attP2 insertion targeting Atx2 CDS 474-494) TRiP.HMS01392 | Bloomington Stock Centre (BDSC) | RRID:BDSC_36114 |
| *Drosophila melanogaster* elavP-gal4 | Bloomington Stock Centre (BDSC) | RRID:BDSC_8760 |
| *Drosophila melanogaster* ppk-CD4-tdTomato | Bloomington Stock Centre (BDSC) | RRID:BDSC_35844 |
| *Drosophila melanogaster* UASt-mCherry.mito.OMM | Bloomington Stock Centre (BDSC) | RRID:BDSC_66532 |
| *Drosophila melanogaster* Ubi-EB1-GFP | S. Rogers (Shimada et al., 2006) |  |
| *Drosophila melanogaster* D42 gal4>UAS ANF GFP | A. Lim, W. Saxton (Lim et al., 2017) |  |
| *Drosophila melanogaster* Nrv2-GFP | Yale FlyTrap database | ZCL2903 |
| Software and Algorithms |  |  |
| Fiji |  |  |
| Graphpad Prism |  |  |
| Nikon Elements |  |  |
| PANTHER GO |  |  |
| Recombinant DNA | | |
| pMT-EOS-alpha-Tubulin | This lab (Barlan et al., 2013). |  |
