## supplementary material for "Ataxin-2 is essential for cytoskeletal dynamics and neurodevelopment in *Drosophila*"

**Supplementary Items**

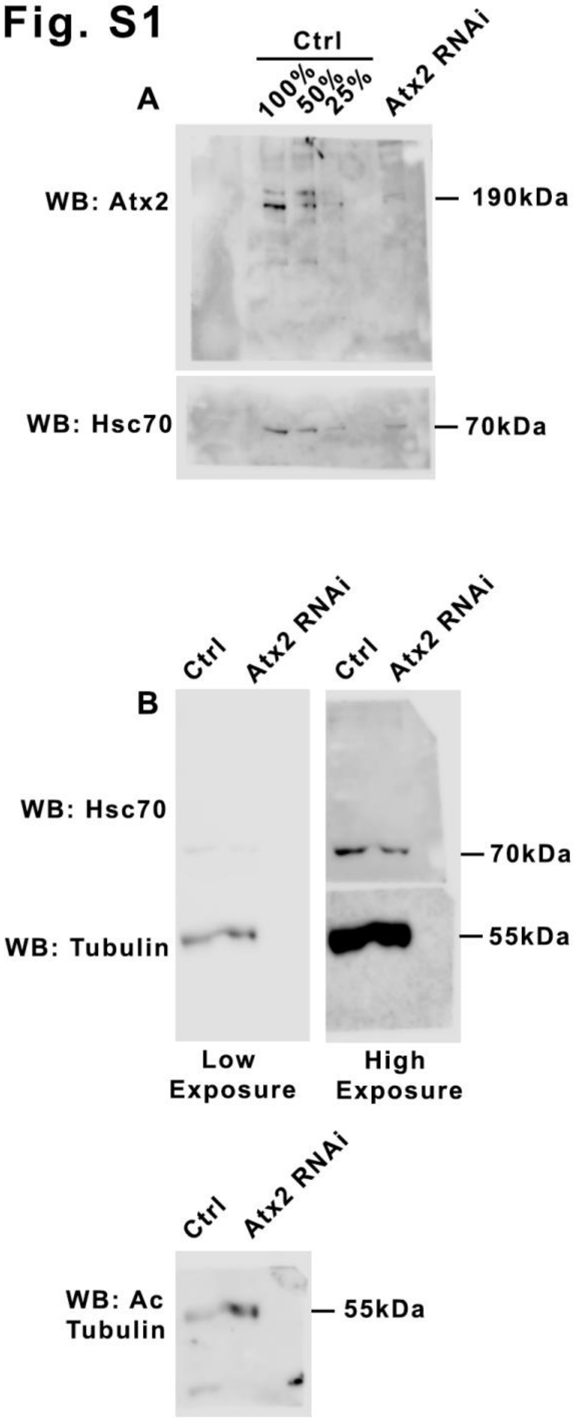


Fig. S1: A. Ataxin-2 depletion in brains by western blot Related to Fig. 1E.
Western blot demonstrating knockdown of Atx2 by elav>Atx2 shRNA #44012. B. Complete western blot membranes showing Acetylated tubulin, total tubulin and Hsc70 (loading control) levels.

**Fig. S2**

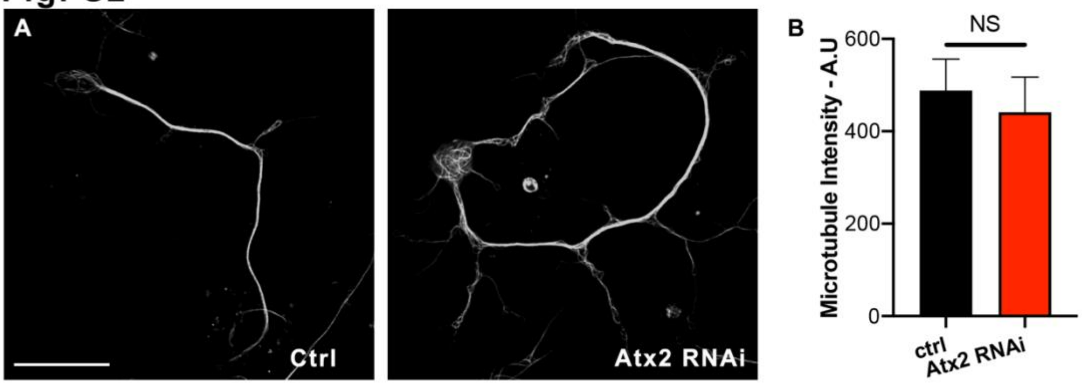


Fig. S2: Microtubule content is equal in control vs Atx2 RNAi neurons in culture. Related to Fig. 1N.

A. Example images showing tubulin staining in extracted neurons. Scale bar, 20µm. B. Quantification of tubulin signal intensity (Control = 488.4 ± 67.7, n = 8 cells, Atx2 RNAi = 441.2 ± 76.1, n = 6 cells, NS p = 0.85).

**Figure S3**

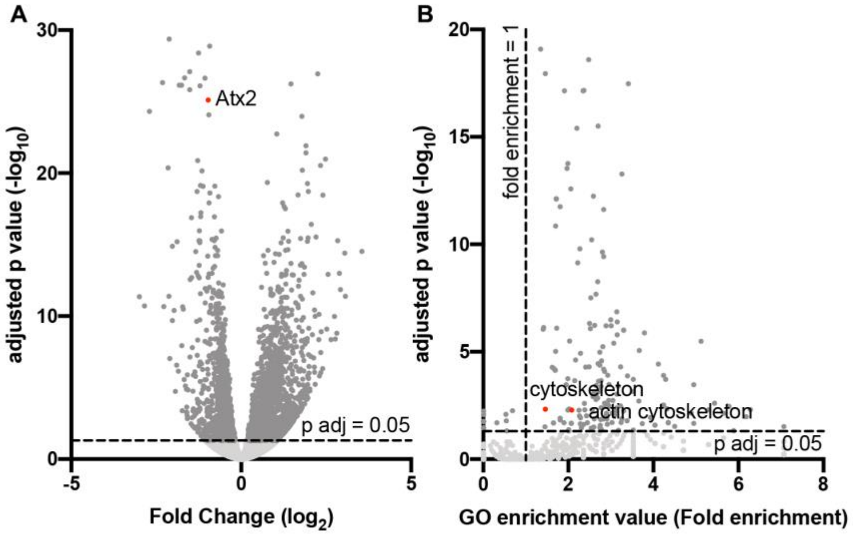


Figure S3. RNAseq analysis in control vs. elav>Atx2RNAi larvae brains shows differential expression of multiple cytoskeletal proteins. Related to Figure 3.

A. Volcano plot showing magnitude of change vs statistical significance of all differentially expressed genes in control vs Atx2 knockdown brains. B. Representation of Gene Ontology analysis cellular component of significantly differentially expressed genes. Fold enrichment compared to reference transcriptome.

Fig S4

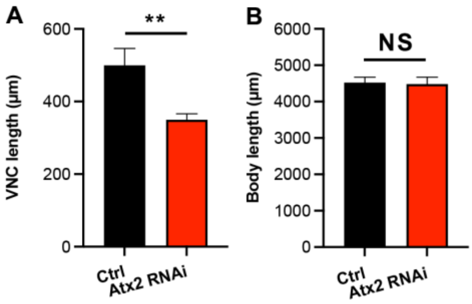


Figure S4. Related to figure 4A-C. A. Length of VNC of Control and Atx2 RNAi animals (Control = 500µm ± 46.7, Atx2 RNAi = 350µm ± 16.5, n = 8 Control and 10 Atx2 RNAi animals, p = 0.0035). B. Length of body of Control and Atx2 RNAi animals (Control = 4523µm ± 152, Atx2 RNAi = 4480µm ± 180, n = 8 Control and 10 Atx2 RNAi animals, NS = not significant).

Table S1: Cytoskeletal genes differentially expressed in elav>Atx2 RNAi brains.

| Protein Class (GO term) | Gene | $Log_2$ FC | FDR adj pvalue |
|---|---|---|---|
| microtubule binding motor protein(PC00156) | Klc | -0.2762914 | 0.0023462 |
| | Kap3 | -0.288398 | 0.02448984 |
| | Kif3C | 1.14466294 | 9.05E-05 |
| | Dhc64C | -0.3329616 | 0.04060265 |
| | Klp3A | -0.3331195 | 0.01395657 |
| | Khc | -0.2767442 | 0.00498595 |
| microtubule or microtubule-binding cytoskeletal protein(PC00157) | CG9313 | 0.84112939 | 0.03499158 |
| non-motor microtubule binding protein(PC00166) | Kat60 | -0.3337638 | 0.00808252 |
| tubulin(PC00228) | betaTub56D | -0.2925545 | 0.03785787 |
| | alphaTub84B | -0.3704931 | 0.0050767 |
| actin binding motor protein(PC00040) | Myo31DF | 0.32221979 | 0.0001427 |
| | up | 1.16374185 | 0.02387464 |
| actin and actin related protein(PC00039) | Arp2 | -0.2474226 | 0.0427629 |
| actin or actin-binding cytoskeletal protein(PC00041) | Zasp52 | 0.39860413 | 0.03571863 |
| | Arpc1 | 0.27096692 | 0.04813796 |
| non-motor actin binding protein(PC00165) | Abp1 | -0.2401069 | 0.04263514 |
| | scra | -0.2459263 | 0.04411634 |

| | | | |
|---|---|---|---|
| | flr | -0.3226018 | 0.0096699 |
| | Fim | -0.5251629 | 0.00108723 |
| | pod1 | -0.9831411 | 6.31E-15 |
| | hts | -0.4116757 | 0.00020457 |
| | cpa | -0.355635 | 0.0011015 |
| | tsr | -0.3179894 | 0.04207127 |
| | SCAR | -0.3788852 | 0.00018631 |
| | Chd64 | 0.88079224 | 3.38E-09 |
| | alpha-Cat | -0.3769759 | 0.01529481 |
| | sn | -0.974603 | 1.16E-06 |
| | Gel | 0.96060323 | 7.11E-07 |
| cytoskeletal protein(PC00085) | Septin 1 | -0.3453684 | 0.03787749 |
| | pnut | -0.3280229 | 0.00016569 |
| | Septin 2 | -0.434462 | 7.82E-05 |
| intermediate filament binding protein(PC00130) | p120ctn | -0.4699921 | 0.00556039 |
| | shot | 0.34320575 | 0.03550769 |
